# A rugged binding landscape unifies static and dynamic paradigms in protein-protein interactions

**DOI:** 10.64898/2026.03.29.715155

**Authors:** Te Liu, Sichao Huang, Wenjian Li, Peiying Wang, Jiahuan Song, Jiaxin Liu, Manjie Zhang, Bin Sun

## Abstract

Predicting protein–protein interaction (PPI) affinities from structural data remains a challenge. Although binding funnel theory describes the formation of native complexes, the topography of the funnel bottom and its influence on affinity are often overlooked. Using two well-controlled nanobody-antigen datasets as model systems, we demonstrate that PPI can exist in both static and dynamic binding paradigms. These two nanobody series adopt nearly identical binding poses toward their respective antigens yet exhibit diverse affinities, each representing a distinct paradigm: a static paradigm in which affinity can be ranked by Rosetta scoring of co-crystal structures alone, and a dynamic paradigm in which affinity can only be ranked using molecular dynamics–sampled ensembles. The two paradigms differ in interfacial dynamics. In the dynamic series, the relative motion between binding partners (Δ*RMSF*) is temperature-sensitive, with optimal affinity correlation at 298 K. In the static series, Δ*RMSF* is minimal and insensitive to temperature. Local frustration analysis establishes a mechanistic bridge between interfacial dynamics and landscape topology. Dynamic interfaces exhibit increased frustration upon thermal sampling, facilitating sampling of functionally relevant microstates, whereas static interfaces show minimal frustration changes across temperatures. The temperature-dependent frustration difference mirrors Δ*RMSF* sensitivity, confirming local frustration as a determinant of interfacial dynamics. Furthermore, static interfaces are characterized by a higher density of canonical hotspot residues, while dynamic interfaces utilize interfacial ruggedness to modulate affinity. Together, these results demonstrate that conserved binding modes can encode different energy landscapes and provide a practical framework for determining when ensemble-based sampling is required for accurate affinity prediction.

## 1 Introduction

Protein-protein interactions (PPIs) orchestrate fundamental biological processes, where the precision of binding and unbinding events determines cellular fate. Central to these interactions is binding affinity, a thermodynamic property traditionally understood through the framework of binding funnel theory [1, 2, 3]. This theory posits that the energy landscape of a PPI is funnel-shaped, guiding two proteins from a vast ensemble of non-specific states toward a native-like global minimum at the funnel bottom. While structural biology has long relied on static complementarity to explain this minimum, whether the funnel bottom is smooth or rugged, and how its topology determines binding affinity, remains poorly understood [4, 5].

Nanobody-antigen interactions provide an ideal model system to investigate these questions. Derived from camelid heavy-chain antibodies, nanobodies feature a highly conserved beta-sheet framework and three hypervariable complementarity-determining regions (CDRs) [6] (Fig. 1A). Their small size (*∼*15 kDa) and simple architecture reduce molecular complexity while retaining the essential biophysics of molecular recognition. Despite this simplicity, nanobodies achieve remarkable functional diversity, recognizing epitopes ranging from flat surfaces to deep catalytic clefts [7] with affinities spanning picomolar to micromolar concentrations (Fig. 1B). Early thermodynamic studies suggested that static, enthalpic contributions from the CDRs (particularly the long, variable CDR3) dominate binding, treating the framework as a passive scaffold [8]. However, recent evidence indicates that conformational dynamics also contribute to nanobody-antigen affinity through entropic effects. For instance, nanobodies targeting the same antigen can achieve high affinity through either enthalpy-driven or entropy-driven mechanisms [9]. Despite their relatively simple fold, the capacity of nanobodies to employ distinct thermodynamic strategies renders them ideal models for studying PPI binding landscapes.

**Figure 1:**
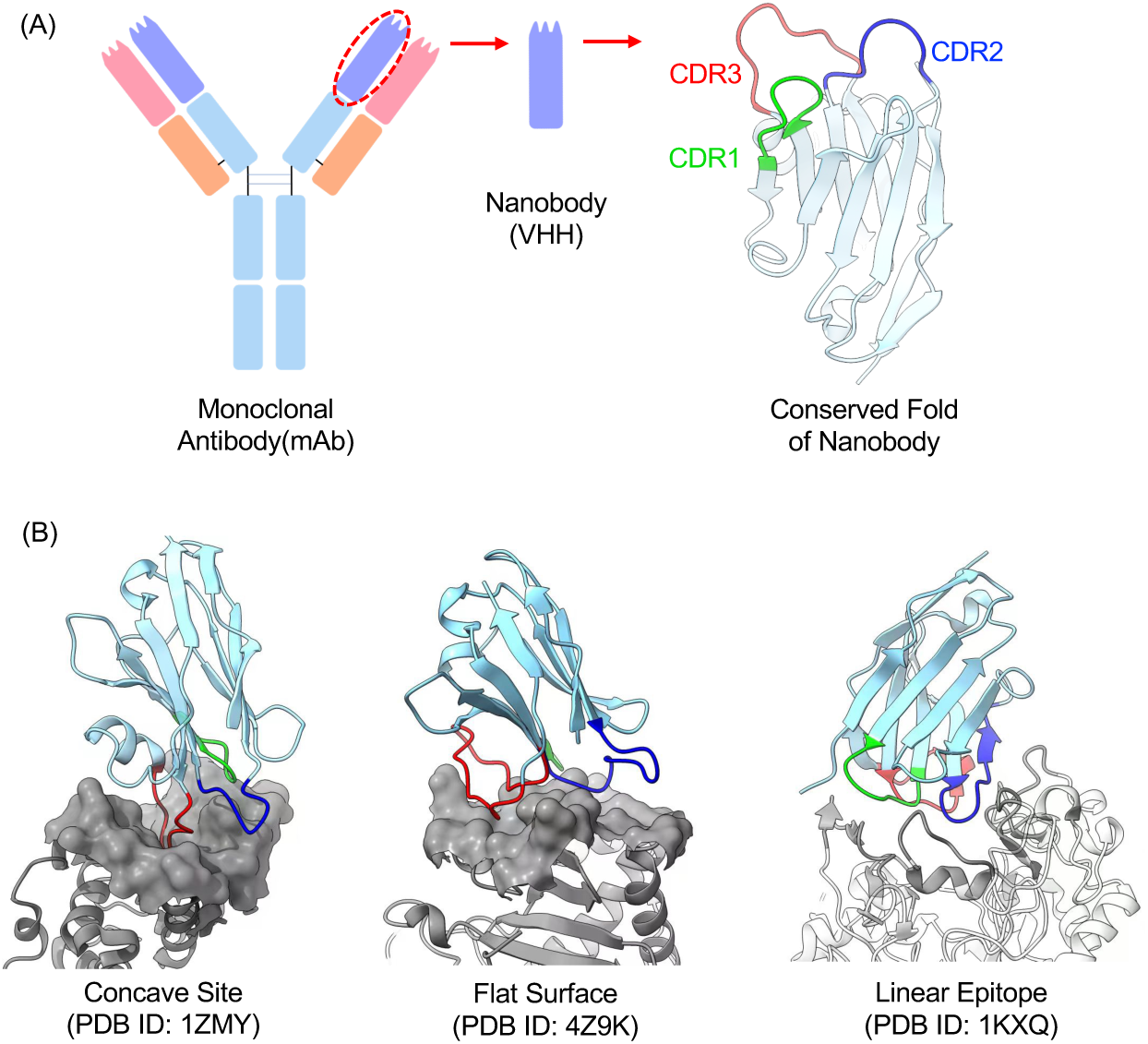
Structural background of nanobody mediated PPI. (A) Schematic illustration of a nanobody, showing its small size relative to a monoclonal antibody and its simple, conserved architecture comprising a highly conserved framework and three CDR loops. (B) Representative nanobody-antigen co-crystal structures demonstrating the nanobody’s ability to recognize diverse epitopes on their cognate antigens.

The growing availability of high-quality co-crystal structures of nanobody-antigen complexes, together with experimentally measured binding affinities [10], now provides an unprecedented opportunity to probe the topography of the binding funnel bottom. These structures, while not necessarily representing the global minimum, are presumed to reside near the bottom of the binding funnel. By systematically evaluating whether such high-resolution structures can accurately reflect experimental affinities, the shape of the underlying energy landscape can be inferred.

In this study, we identified two nanobody series from reported nanobody-antigen studies [10], each containing multiple nanobodies that exhibit near-identical binding modes toward their respective antigen yet display diverse affinities. These two sets revealed fundamentally different binding mechanisms: one where affinity is accurately predictable from static co-crystal structures, and another governed by interfacial dynamics. This finding establishes interface dynamics as a critical determinant of affinity in dynamic binding paradigms. We further demonstrate that only at an appropriate temperature around 298 K do interfacial dynamics sample the appropriate microstates to reflect experimental affinity. Using local frustration theory, we relate these dynamical differences to the degree of funnel bottom ruggedness, showing that PPI binding landscapes can be either smooth or rugged even when co-crystal structures exhibit convergent binding modes. Together, our work unifies these observations into a single framework, identifying PPI interfacial dynamics as the missing link that explains why universal affinity prediction has remained challenging relying on purely structural data.

## 2 Methods

### 2.1 Nanobody-antigen structural and affinity data collection

We collected all reported nanobody-antigen co-crystal structures with experimental affinity measurements from the SAbDab-nano database [10] (accessed December 1, 2023). Additionally, we surveyed the literature to identify 10 data points [11] not included in the SAbDab-nano database, resulting in a total of 95 nanobody-antigen complexes.

For the 10 additional data points from [11], all nanobodies bind to the same epitope of the SARS-CoV-2 spike protein with different affinities. Among these, five nanobodies have co-crystal structures resolved with corresponding PDB entries deposited in the Protein Data Bank, while the remaining five are designed mutants derived from the same parent nanobody (PDB 7Z1A). For these five nanobodies (detailed in Section 3.2), only affinity measurements were available; their complexes with the antigen were modeled using the parent structure PDB 7Z1A via SWISS-MODEL [12]. We then applied the Rosetta side-chain relaxation protocol, a well-vetted tool for fine-tuning side-chain conformations [13, 14], to further optimize side-chain packing after homology modeling. Although this modeling strategy bears limitations owing to its in silico nature, it is likely to provide reasonable starting structures. First, the highly conserved binding mode of the 7Z1X series greatly reduces the search space. Second, RMSD analysis from 100 ns MD simulations shows that the modeled structures are comparably stable to co-crystal structures (Fig. S1), indicating that Rosetta side-chain optimization was effective.

From the total 95 nanobody-antigen complexes, we further processed the PDB files by removing entries with incorrect affinities and redundant structural information, yielding 83 structures for analysis. For these 83 structures, we first applied the Rosetta re-docking protocol to assess whether if there exists a binding funnel and if each complex resides near the bottom of a binding funnel. We identified 78 complexes that satisfy this criterion and 5 that do not. All nanobodies, along with their structural information (PDB IDs), experimental affinities (including assay type and measurement temperature), and funnel-bottom classification, are listed in Table S1. The nanobody-antigen affinity data were originally reported as dissociation constants (*K_d_*). For our usage, we converted *K_d_* values to binding free energies (Δ*G_exp_*) via:

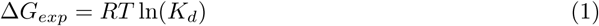

where *R* is the gas constant, and *T* is the temperature at which the *K_d_*values were measured.

### 2.2 Rosetta re-docking

We used the Rosetta docking protocol (RosettaDock) [15] to redock each nanobody to its antigen, assessing whether a binding funnel exists and whether the co-crystal structures reside near its bottom. If such a funnel is present, decoy conformations that closely resemble the native nanobody-antigen binding mode should exhibit lower binding free energies. We first applied *relax.linuxgccrelease* module [16] to optimize side chains and minimize energy, eliminating steric clashes and unfavorable interactions within the structures. Following relaxation, we used *docking prepack protocol.linuxgccrelease* module to ensure that side chains outside the docking interface adopt low-energy conformations, a prerequisite for accurate decoy scoring. Finally, we generated 1000 docking decoys for each complex using *docking protocol.linuxgccrelease*. The interface energy (scored via the REF 2015 scoring function [17]) and interface RMSD relative to the co-crystal structure were calculated to evaluate decoy quality and assess funnel characteristics.

### 2.3 Computational evaluation of nanobody-antigen binding affinities from co-crystal structures using multiple methods

For nanobody-antigen complexes identified as possessing funneled binding landscapes, we employed multiple computational methods to calculate binding affinities directly from the co-crystal structures. These methods include FireDock [18], Bluues [19], ZDock [20], IRAD [21], and the Rosetta REF15 scoring function [17]. The first four methods have been previously reported to predict antigen-antibody binding affinities [22], while Rosetta REF15, the default scoring function within the Rosetta software framework, has become a standard energy function in structural biology applications Our aim was to investigate whether the reported nanobody-antigen co-crystal structures can recapitulate experimental affinities and whether the results exhibit method dependence. For FireDock, Bluues, ZDock, and IRAD, we utilized the PARCE open-source framework [23], which integrates these methods into a single platform. Affinity calculations were performed using the crystal structures as input with default scoring parameters. For Rosetta scoring, affinities were evaluated by running the locally installed *score jd2.linuxgccrelease* executable.

### 2.4 Molecular dynamics simulations

All conventional molecular dynamics (MD) simulations were performed using the AMBER20 package [24]. Each nanobody-antigen complex in the 2P4X and 7Z1X series was solvated in a 12 Å margin water box with 0.15 M KCl ionic strength using the TLEAP program. The AMBER ff19SB force field [25] and the OPC water model [26] were used for all systems. Each system was first subjected to 50,000 steps of energy minimization, with the first 200 steps using the steepest descent algorithm and the remaining steps using the conjugate gradient algorithm. The minimized system was then heated from 0 K to 300 K via a two-step procedure: 0 to 100 K in the NVT ensemble over 0.1 ns, followed by 100 to 300 K in the NPT ensemble over 0.5 ns. Harmonic constraints with a force constant of 5 kcal/mol/Å^2^ were applied to protein backbone atoms during heating. After heating, a 1 ns equilibrium simulation was performed at 300 K with a reduced force constant of 1 kcal/mol/Å^2^. Finally, 100 ns production simulations were initiated from the equilibrated system in the NPT ensemble at 300 K. Our choice of running relatively short 100 ns production simulations is rationalized by the fact that these co-crystal structures reside near a funnel bottom (Fig. S2), where large-scale structural reorganization is energetically unfavorable. Thus, 100 ns simulations should be sufficient to sample equilibrated conformations around the local minimum. This approach is further supported by numerous studies demonstrating that longer MD simulations do not necessarily yield more accurate Δ*G* values relative to experimental data, as repeatedly reported in MM-GBSA/PBSA studies [27, 28].

The SHAKE algorithm [29] was used to constrain all bond lengths involving hydrogen atoms, and the Langevin thermostat was employed for temperature control. Long-range electrostatic interactions were handled using the Particle-Mesh Ewald (PME) method [30]. The time step was 4 fs after hydrogen mass repartitioning [31], and the nonbonded cutoff was 12.0 Å. For each nanobody-antigen complex, 100 ns production MD simulations of the isolated nanobody and antigen (starting from the structure extracted from the complex) were also performed. In addition to simulations performed at 300 K, we repeated MD simulations for each complex in the 2P4X and 7Z1X series at temperatures ranging from 285 K to 315 K in 5 K increments. These simulations were carried out for both the nanobody-antigen complexes and the corresponding isolated nanobodies and antigens.

To assess whether our 100 ns sampling is sufficient for convergence—and given that our final goal is to use these trajectories for Rosetta-based affinity calculations—we repeated the 100 ns simulations in triplicate for all nanobody-antigen complexes at 295 K, 300 K, and 305 K. As we show in the Results section, individual replicas and the average of the triplicate runs at these three critical temperature points yield consistent energy scoring. Furthermore, structural deviation analysis using heavy-atom RMSD relative to the starting co-crystal structure indicates that, for both the 2P4X and 7Z1X series across the 295 K–305 K range, the nanobody-antigen complexes underwent limited structural reorganization within 100 ns, characterized by RMSD values of approximately 2–2.5 Å (see Fig. S1). A summary of all MD simulations performed in this study is provided in Table S2.

### 2.5 Rosetta-based scoring of the MD-sampled nanobody-antigen ensembles

In the present study, we focus on affinity ranking within each nanobody series rather than absolute binding free energy prediction. Our goal is to compare whether scoring a single co-crystal structure versus scoring multiple MD-sampled structures better reflects the experimental affinity trend. We elected to use the Rosetta REF2015 scoring function [17], a widely used tool with proven accuracy, to score various nanobody-antigen complex structures and estimate the Δ*G* reflected by these structures. This ranking-based approach alleviates the stringent sampling requirements of rigorous free energy calculations while remaining sufficient for our purpose.

Frames were extracted from the 100 ns production MD trajectories of each nanobody-antigen complex at 1 ns intervals, yielding 100 frames per trajectory for subsequent Rosetta scoring. Each frame, with waters and ions removed, was used as input to the score jd2.linuxgccrelease executable to score the nanobody-antigen complex (*G*_com_), the isolated nanobody (*G*_vhh_), and the isolated antigen (*G*_antigen_). The calculated Δ*G* for each frame was defined as:

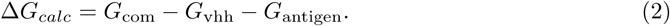

The final Δ*G_calc_*value for each complex was reported as the average over the 100 frames. Dry protein conformations extracted from MD trajectories sampled at temperatures ranging from 285 K to 315 K were all subjected to Rosetta scoring. This approach of feeding only dry protein structures into Rosetta has an additional advantage for our study. Because REF2015 uses fixed energy terms, including solvation, to score a given protein conformation [17], the scoring function reflects only what is encoded in the protein structures themselves. This feature is actually advantageous for our purpose of exploring how conformational dynamics affect binding affinity, as it allows us to isolate the structural contributions to Δ*G* without confounding temperature-dependent solvation effects.

One potential caveat of using only dry structures is the omission of solvation effects, which could introduce bias into the calculated Δ*G* values. However, our solvation energy calculations on these dry protein structures at different temperatures show that the solvation free energy for both the 2P4X and 7Z1X series remains largely stable across the 285 K–315 K temperature range (see details in Fig. S3). This stability further justifies our use of REF2015 to score the complexes.

### 2.6 Local frustration analysis

We performed local frustration analysis using frustratometeR [32], an R package that implements the frustratometer algorithm with additional functionalities for analyzing molecular dynamics trajectories. Specifically, we calculated configurational frustration for both the co-crystal structures and the MD trajectories of each nanobody-antigen complex. For PDB analysis, each co-crystal structure was used directly as input to frustratometeR with default parameters. For MD analysis, frustratometeR reported average frustration values over all frames contained in each trajectory. We focused on the percentage of highly frustrated interaction pairs within radial shells at varying distances from the nanobody-antigen interface center. Specifically, we calculated the radial distribution function of highly frustrated pairs around the interface center. The interface center was defined as the geometric center of all residues from both binding partners located within 5 Å of each other. For PDB analysis, this interface center was fixed for each structure. For MD analysis, the interface center was determined from the first frame, and all subsequent frames were aligned to this reference frame prior to frustration calculation. For each nanobody series, the PDB-based radial distribution function was averaged across all structures in the series, and the MD-based radial distribution function was similarly averaged across all corresponding trajectories. The difference between the PDB-based and MD-based radial distribution functions was then calculated for each series. Additionally, because we performed MD simulations at multiple temperatures (285 K to 315 K), we investigated the temperature sensitivity of the frustration difference. For each simulation temperature, we calculated the MD-based radial distribution function of highly frustrated pairs from the corresponding trajectory. These were then compared to the PDB-based radial distribution function, which was calculated once from the static co-crystal structure and served as a constant reference across all temperatures.

### 2.7 Hotspot residue analysis

We performed interface hotspot residue analysis using the *Flex ddG* protocol [33] within the Rosetta software suite. This protocol consists of four main steps. First, global minimization optimizes the backbone and side chain torsions of the input structures. Second, the *backrub* protocol [34] is applied to sample local backbone conformational flexibility. Third, the binding free energy (ΔG) is scored for both the wild-type complex and the complex with introduced mutations. Finally, the ΔΔG values (mutant minus wild-type) are calculated.

Because nanobodies within each series bind to their cognate antigen using nearly identical binding modes, they share common binding interfaces. This allowed us to select fixed sets of interfacial residues for alanine scanning within each series. For the 2P4X series, we selected 12 interface residues that form direct contacts between nanobody and antigen: antigen residues N62, T70, Y73, E111 and Y115; and nanobody residues Y151, Y153, I156, G223, Y225, R230, and T231 in 2P42. For the 7Z1X series, we selected 17 interface residues: antigen residues K112, G114, Y117, N118, E152, F158, Q161 and S162; and antigen residues R223, F225, R248, S253, R296, V297, T298, R299, and S300 in 7Z1A. In each nanobody-antigen co-crystal structure, these amino acids were mutated to alanine, and the ΔΔG values were calculated using *Flex ddG*. All code and scripts used for this project are available at a public repository at: https://github.com/bsu233/bslab/tree/main/2026NBfunne

## 3 Results

### 3.1 Most resolved nanobody-antigen co-crystal structures are located near the bottom of a binding funnel

To assess whether experimentally determined nanobody-antigen co-crystal structures follow binding funnel theory, we performed local perturbation and re-docking using the RosettaDock protocol for each resolved complex, generating 1000 decoy structures per system (Fig. 2A). The interface RMSD (iRMS) and interface energy for each decoy were calculated using the Rosetta InterfaceAnalyzer module [35]. A well-formed binding funnel is characterized by a strong correlation between iRMS and interface energy, where the crystal structure, or decoys very close to it, exhibits the most favorable energy.

**Figure 2:**
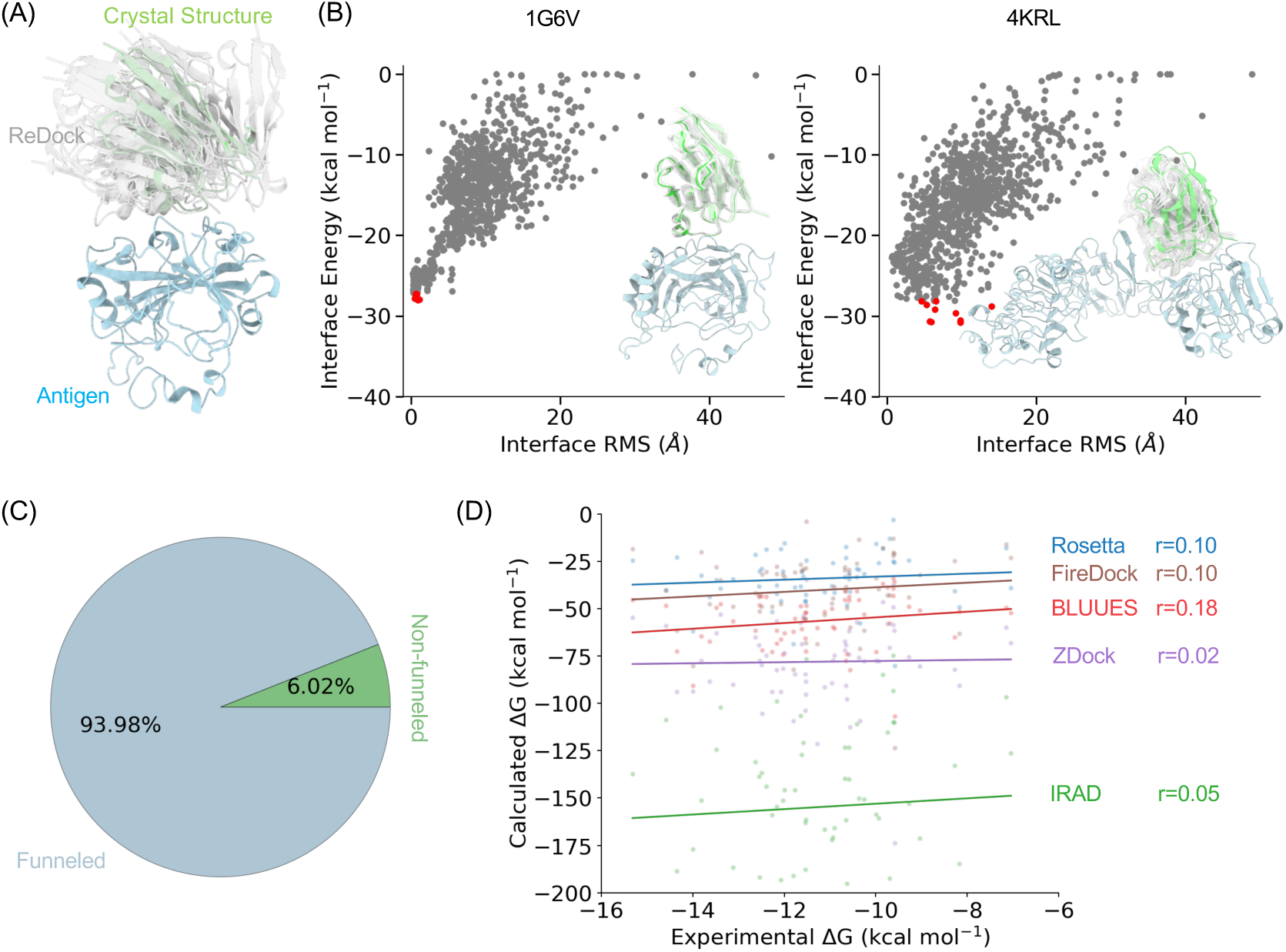
Nanobody-antigen co-crystal structures analysis. (A) Binding funnel evaluations. The nanobody in co-crystals was perturbed and redocked back to the antigen. The decoy’s interface energy was scored using Rosetta REF15 scoring function [17] and was plotted against its deviation from co-crystal binding mode measured by interface RMS. (B) A typical binding funnel seen in PDB 1G6V [37] and a non-funnel binding in PDB 4KRL was shown. The red dots represent the 10 decoys with the lowest Rosetta-scored interface energies. (C) The percentage of funneled and non-funneled binding in the collected 83 nanobody-antigen co-crystal structures. The detailed redocking results for each co-crystal is shown in Fig. S2. (D) For all 78 funneled complexes, the correlation between calculated ΔGs with experimental affinities using various scoring functions were plotted.

Fig. 2B shows representative examples of nanobody-antigen binding that exhibit funneled and non-funneled behavior, respectively. For the funneled binding (PDB 1G6V), the structural overlay of the ten lowest-energy decoys demonstrates tight clustering around the crystal structure. The corresponding energy landscape shows a clear funnel, with the lowest-energy decoy achieving an iRMS of 0.531 Å and an interface energy of *−*28.013 kcal/mol. In contrast, for a non-funneled binding such as PDB 4KRL [36], the lowest-energy decoy structures span a wide range of iRMS values, in contrast to the tight clustering observed around the native binding mode in funneled systems (Fig. 2B). Among the 83 nanobody-antigen co-crystals we collected, 78 exhibit funneled landscapes resembling PDB 1G6V, while 5 exhibit non-funneled landscapes with poorly defined energy minima (detailed re-docking results are shown in Fig. S2).

We next focused on these funneled complexes to investigate whether binding affinities could be directly predicted from static co-crystal structures. Nanobodies in these funneled structures exhibit a highly conserved framework, with pairwise RMSD of the *β*-sheet framework below 2.0 Å and RMSF of the framework among crystal structures around 0.5 Å (Fig. S4). We computed binding free energies (ΔG) using a panel of established computational methods, including the Rosetta REF15 score function, IRAD, Bluues, ZDock, and FireDock. Interestingly, none of these methods yielded a significant correlation with experimentally measured affinities across the entire dataset, with correlation coefficients all below 0.2 (Rosetta: r = 0.10, FireDock: r = 0.10, Bluues: r = 0.18, ZDock: r = 0.02, and IRAD: r = 0.05; see Fig. 2D). This failure persisted even when complexes were grouped by the experimental technique used for affinity measurement (e.g., ITC vs. SPR; Fig. S5), suggesting that methodological differences alone could not explain the poor performance. Therefore, while most resolved nanobody-antigen co-crystal structures reside near the bottom of a binding funnel, static structure-based scoring across this diverse set yields poor correlation with experimental affinities, prompting us to investigate system-specific factors that may govern affinity prediction.

### 3.2 Identifying static and dynamic binding paradigms in two nanobody sets, respectively

Since the 78 nanobody-antigen data were collected by different labs where multiple variables such as antigen, epitope, and affinity measurement method were inconsistent. We therefore constructed curated sub-datasets that have the nanobody-antigen structures and affinities determined by the same research group to enable reliable comparison. This filtering yielded two high-quality datasets: the 2P4X series from Tereshko *et al* that contain nine nanobodies bind to the RNase A protein [38], and the 7Z1X series from Mikolajek *et al* that contain ten nanobodies bind to the receptor binding domain (RBD) of the spike protein of SARS-COV-2 [11] (Fig. 3). The nanobodies within each set bind to the same epitope on their respective antigen (pair-wise complex RMSD values *<*2 Å), yet they exhibit experimental affinity difference up to 5 kcal/mol (Fig. 3). Rosetta-calculated ΔG shows that, in the 2P4X series, the static co-crystal structures are sufficient to rank the experimental affinity, with co-efficient r=0.80. In contrast, for the 7Z1X series, the static-structure based co-efficient r=*−*0.41, demonstrating that the former set relies on a static co-crystal structure, and the latter does not.

**Figure 3:**
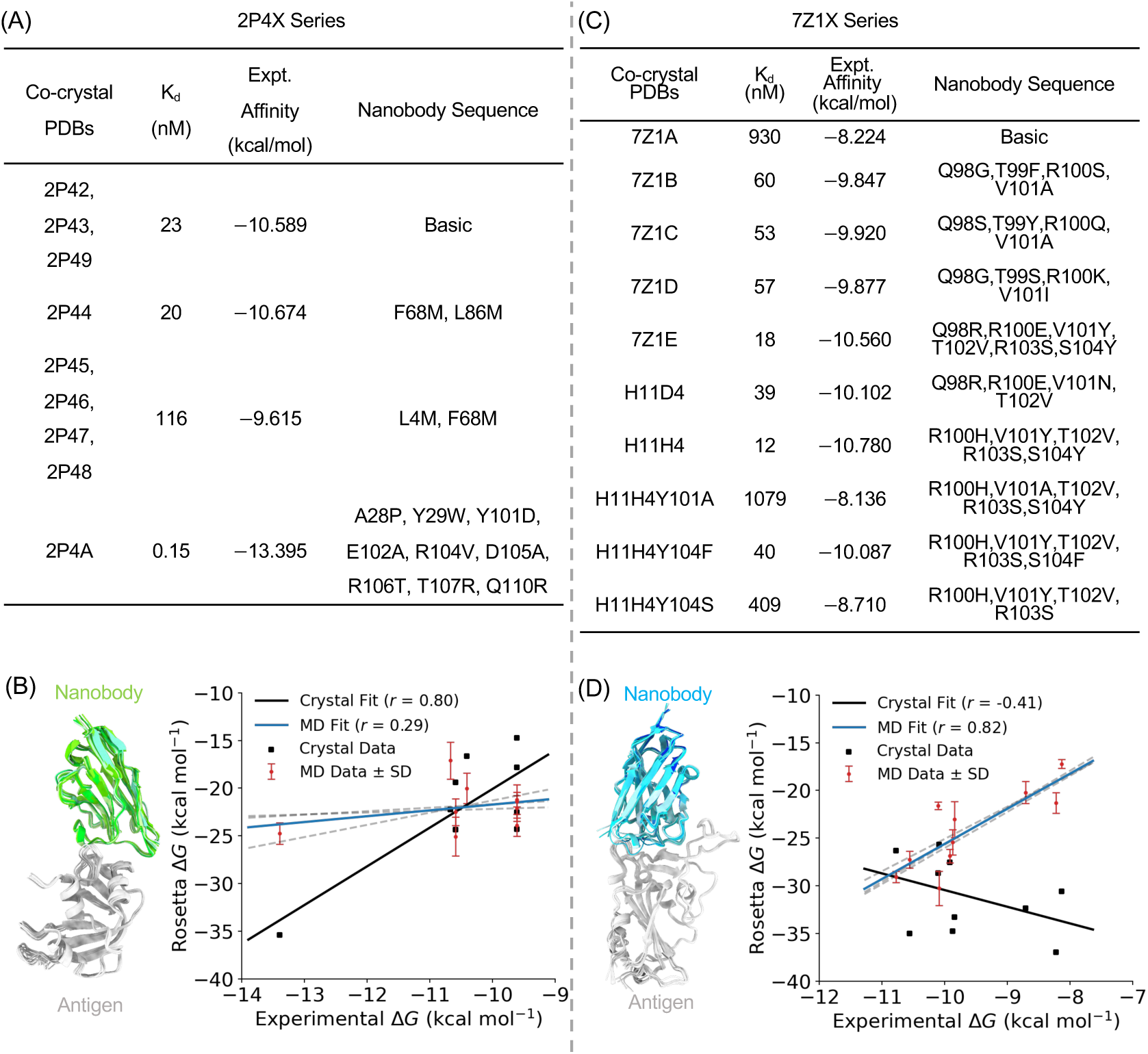
Two nanobody datasets with nanobodies binding to the same epitope on the antigen yet exhibiting diverse affinities. (A) PDB IDs of nanobody-antigen co-crystal structures in the 2P4X series [38], experimentally measured dissociation constants (*K_d_*), binding free energies (in kcal/mol) converted from *K_d_* via Eq. 1, and nanobody sequence variations relative to the first structure listed (PDB 2P42). (B) Overlay of all nine co-crystal structures aligned on the antigen. Correlation of Rosetta-scored Δ*G* values based on co-crystal structures and on 100 ns MD-sampled conformations (at 300 K) with experimental affinities. Three independent MD runs were performed; the average Rosetta Δ*G* over the three runs is fitted against experimental values The blue line represents the average value over three MD runs, and the gray dashed lines represent the individual MD values. Correlations from individual MD runs are shown in Fig. S6. (C-D) Data for the 7Z1X series [11]. Notably, the last five nanobodies lack resolved co-crystal structures; their nanobody-antigen complex structures were modeled based on PDB 7Z1A.

To probe the extent to which conformational dynamics affect the affinity ranking, especially for the 7Z1X series, we preformed 100 ns production MD simulations of the nanobody-antigen complex, and calculated the Rosetta ΔG using the MD sampled conformations. Remarkably, the 7Z1X series now exhibited excellent correlation with experimental affinities, with r=0.82. Meanwhile, the MD-ensemble based ΔG for the 2P4X series became greatly worsened, leading r=0.29. This suggests that for the 2P4X series, the nanobody-antigen affinities are encoded in the crystal structures, and further averaging over conformational dynamics adds noise rather than insight. While the 7Z1X series exhibited the opposite pattern, and their affinities are not determined by a single conformation but is an property of the dynamic ensemble. Thus, our analysis of curated datasets reveals that nanobody-antigen binding exists two distinct binding paradigms, a static paradigm represented by 2P4X series, and a dynamic paradigm represented by the 7Z1X series.

### 3.3 The dynamics of binding interface as a determinant of affinity in the dynamic binding paradigm

While 7Z1X series requires MD ensembles for accurate affinity prediction and 2P4X series does not, suggesting that binding free energy for the former is not localized to a single structural minimum but is instead distributed across a conformational ensemble. We next investigated the temperature sensitivity of these ensembles in reporting affinities (Fig. 4A). The 7Z1X series exhibits clear sensitivity to simulation temperature, as increasing the temperature from 285 K to 315 K results in a correlation coefficient that first increases, then decreases, with optimal correlation achieved near room temperatures of 295 K (r = 0.90) and 300 K (r = 0.81). In contrast, the 2P4X series remains poorly correlated and largely insensitive to temperature across the 285–315 K range, with the minor variations representing stochastic noise rather than meaningful temperature dependence. To assess fitting robustness, two additional 100 ns MD replicas were performed at three critical temperatures (295 K, 300 K, and 305 K), which yielded consistent results (Fig. 4A).

**Figure 4:**
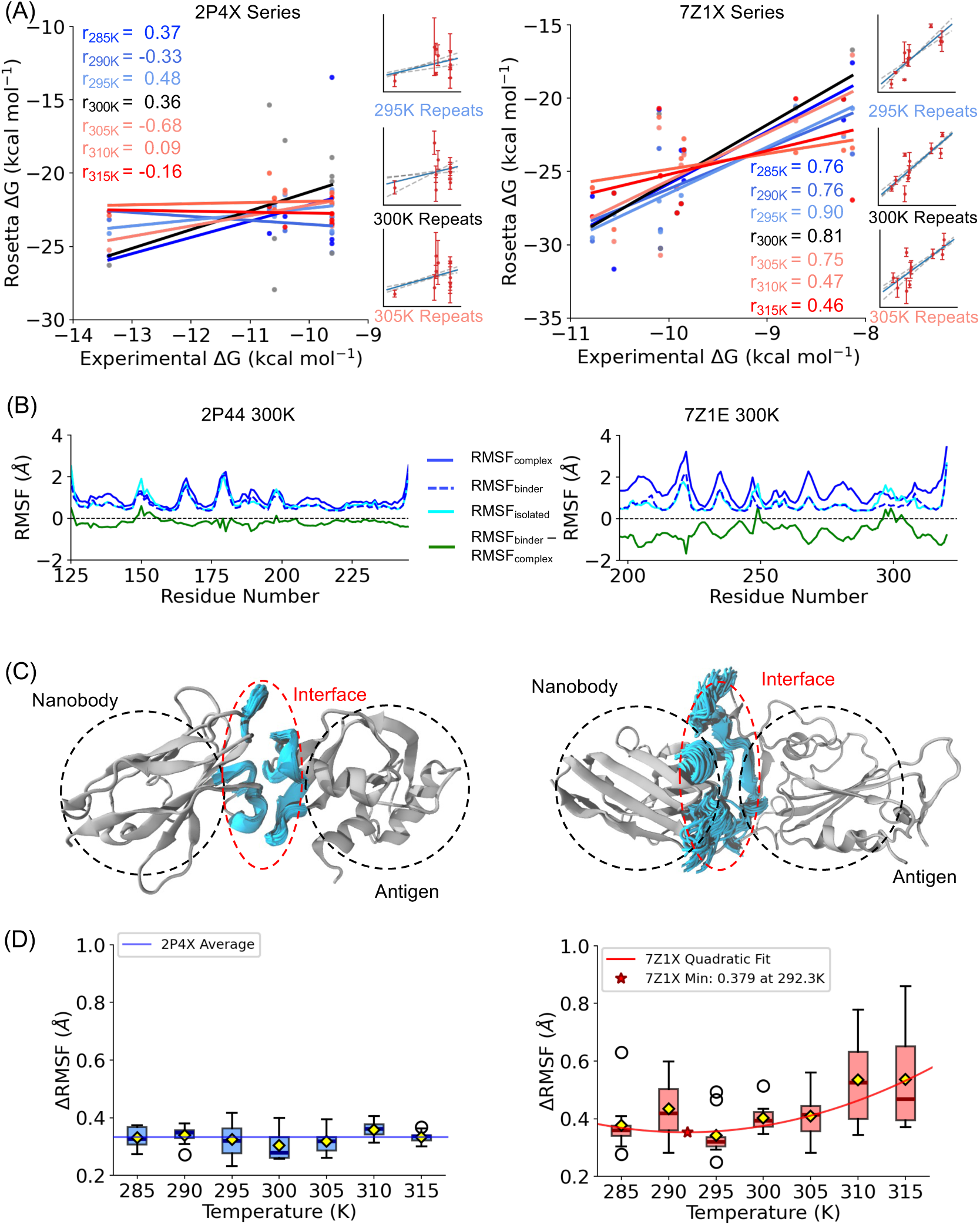
Temperature sensitivity of binding interface dynamics. (A) Correlation between Rosetta-calculated Δ*G* (using 100 ns MD-sampled ensembles at different temperatures) and experimental affinities. At three critical temperatures (295 K, 300 K, and 305 K), two additional 100 ns MD replicas were performed, and the average Δ*G* from these runs was also fitted against experimental affinity (see Fig. S9 for more details). (B) *RMSF_complex_*, *RMSF_binder_*, and *RMSF_isolated_* for two representative data points from the two nanobody series at 300 K. The *RMSF_binder_ − RMSF_complex_* difference is shown as a green curve. RMSF analysis for all other data points is shown in Fig. S8. (C) Superimposition of MD-sampled conformations aligned onto the complex for PDB 2P44 and 7Z1E. (D) Temperature-dependent Δ*F* for the two nanobody series. Δ*RMSF* values for the 2P4X series were fitted to a constant value (Δ*RMSF* = 0.329 Å), while those for the 7Z1X series were fitted to a quadratic curve.

The nonmonotonic temperature dependence of the correlation coefficient for the 7Z1X series likely arises from two physical effects. At lower temperatures (below 290 K), the system may suffer from broken ergodicity, where the equilibrium distribution of functionally relevant microstates is inadequately sampled. At higher temperatures (above 305 K), the complexes sample high-entropy states that deviate from the functional binding interface, introducing noise that obscures the affinity signal. The existence of an optimal correlation near 295–300 K corroborates this explanation. Furthermore, heterogeneity in the temperature sensitivity of individual nanobodies within the 7Z1X series (Fig. S7), because each nanobody differs by multiple residues and represents a distinct PPI entity, also likely contributes to the nonmonotonic behavior at the individual complex level.

To characterize the physical nature of these dynamics, we evaluated residue-wise RMSF from MD trajectories. For each nanobody complex, three RMSF profiles were calculated (Fig. 4B): *RMSF_complex_* was obtained by aligning trajectories on the entire complex; *RMSF_binder_* was obtained by aligning on the individual nanobody and antigen, respectively; and *RMSF_isolated_* was calculated from simulations of the isolated binders. The difference *RMSF_binder_ − RMSF_complex_* therefore reports on the relative motion between the two binding partners within the complex (Fig. 4B). We observed that internal structural fluctuations of the individual components remain relatively constant upon binding, as *RMSF_binder_* and *RMSF_isolated_* are nearly identical for both datasets (Fig. 4B). However, the *RMSF_binder_ − RMSF_complex_* difference curves, which reflect interfacial dynamics, differ markedly between the two series. For a representative 7Z1X data point (PDB 7Z1E), *RMSF_binder_*values for most residues are smaller than *RMSF_complex_*, yielding difference curves that deviate substantially from zero. This indicates considerable relative motion between the two binding partners during the simulation. In contrast, for a representative 2P4X data point (PDB 2P44), the *RMSF_binder_− RMSF_complex_*differences are near zero, indicating negligible relative motion and thus minimal interfacial dynamics. This pattern holds across all data points in each series (Fig. S8). Superimposing the MD-sampled conformations aligned onto the nanobody-antigen complex clearly shows that, for PDB 7Z1E, the interfacial regions of the complex sample more diverse conformations than those of PDB 2P44 (Fig. 4C).

To quantify interfacial dynamics, we defined a metric 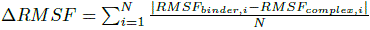 (where *i* indexes residues). The temperature dependence of Δ*RMSF* clearly differentiates the two binding paradigms (Fig. 4D). The 2P4X series exhibits low and nearly invariant Δ*RMSF* (0.329*±*0.016 Å) across all simulation temperatures, consistent with a rigid, lock-and-key interface where thermal energy within the 285–315 K range is insufficient to induce significant relative motion. In contrast, the 7Z1X series shows a higher baseline magnitude of Δ*RMSF* values. Moreover, Δ*RMSF* exhibits a nonmonotonic temperature response: values are minimized near room temperature (295–300 K) and increase at both lower and higher temperatures. Below 290 K, Δ*RMSF* increases slightly, consistent with broken ergodicity where the system lacks sufficient thermal energy to explore the equilibrium distribution. Above 300 K, Δ*RMSF* increases sharply, accompanied by greater variance, indicating that elevated temperatures drive the ensemble into high-entropy states that deviate from the functional binding interface. The temperature-dependent Δ*RMSF* values are well described by a quadratic fit (R^2^ = 0.757), with a theoretical minimum at approximately 292.3 K. Notably, for the 7Z1X series, both the Δ*RMSF* minimum and the optimal affinity correlation converge near 295–300 K. This convergence demonstrates that room temperature (at which the affinities were measured) provides the appropriate thermal energy to sample the functional ensemble. Together, these findings establish that binding interface dynamics are a primary determinant of affinity in the dynamic recognition paradigm, and that accurate computational affinity ranking requires proper sampling of these interfacial degrees of freedom.

### 3.4 Local frustration analysis reveals the topography of the binding funnel bottom

We showed that the two nanobody series exhibit different interfacial dynamics which contribute differentially to binding affinities, we next investigated whether the topography, or ruggedness, of the binding funnel bottom differs between the two series. Given that both series have co-crystal structures residing near the funnel bottom, the different dynamical behaviors and their affinity relevance suggest underlying differences in the energy landscape. To test this hypothesis, we employed local frustration theory to analyze the co-crystal structures and the MD trajectories sampled at different temperatures. Local frustration theory quantifies residual energetic conflicts within a protein structure by comparing the stability of native residue-residue interactions to a decoy ensemble [39]. Residue pairs are classified as minimally frustrated, neutral, or highly frustrated based on whether the native interaction confers favorable or unfavorable thermodynamic stability relative to decoys. The elegance of local frustration theory lies in its ability to retrieve energetic information encoded in static structures that can be translated to protein functional dynamics [39, 40, 41], thus it can serve as a mechanistic bridge connecting our observed interfacial dynamics to the underlying energy landscape that governs nanobody-antigen interactions.

We focused on the distribution of highly frustrated interactions across the PPI interface regions (Fig. 5A). We first used representative data points from each series (PDB 2P44 and PDB 7Z1E) to qualitatively demonstrate the frustration difference between the two nanobody series. For PDB 2P44, MD sampling at 300 K did not alter the percentage of highly frustrated pairs in the interface region compared to PDB-based analysis. In contrast, for PDB 7Z1E, MD sampling introduced a considerable increase in highly frustrated interaction pairs (Fig. 5A). To quantify this observation, we plotted the radial distribution function of the percentage of highly frustrated pairs around the interface center (Fig. 5B), averaged over all nanobody-antigen complexes within each series. For the 7Z1X series, the MD-derived ensembles exhibit a significantly higher percentage of highly frustrated residues at the binding interface (0 to 10 Å shell) compared to the co-crystal structures. In contrast, for the 2P4X series, the MD and PDB frustration profiles are nearly identical within the 0 to 10 Å region.

**Figure 5:**
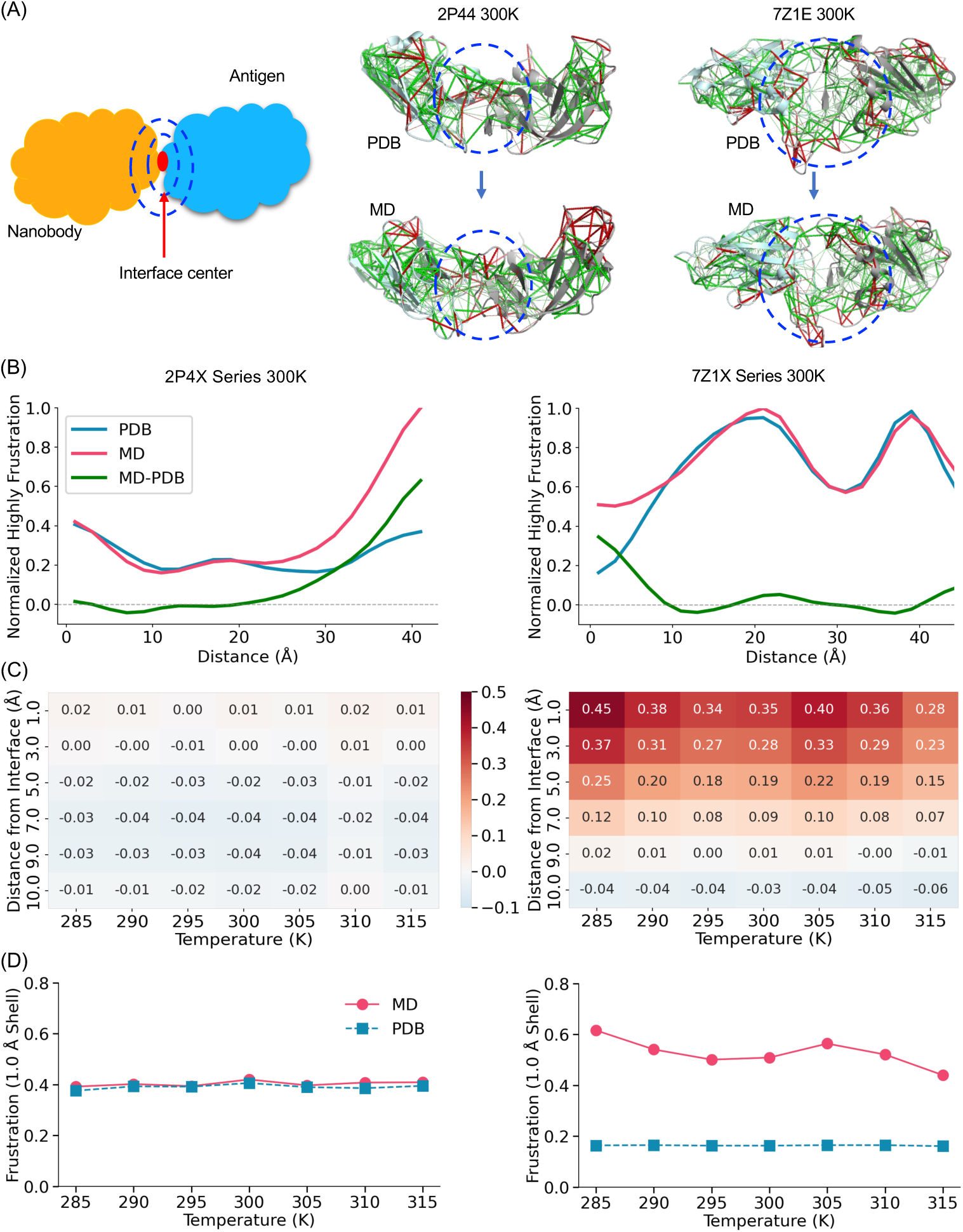
Local frustration patterns around the binding interface. (A) Schematic illustrating the radial distribution function (RDF) of the percentage of highly frustrated interaction pairs across the PPI binding interface. Shown on the right are two representative data points with frustration results based on PDB structures and MD trajectories at 300 K. Minimally frustrated and highly frustrated pairs are indicated by green and red lines, respectively. (B) Radial distribution function of the percentage of highly frustrated interaction pairs. The x-axis represents the distance from the interface center, and the y-axis represents the normalized percentage of highly frustrated residue pairs relative to the total number of contacts in each radial shell. (C) Heatmaps depicting the temperature sensitivity of the frustration difference (MD minus PDB) from 285 K to 315 K within the 0 to 10 Å shell. Detailed data are shown in Fig. S10. (D) Detailed temperature sensitivity of the frustration difference (MD minus PDB) within the 1 Å shell encompassing the interface center.

To investigate the temperature sensitivity of interfacial frustration patterns, we focused on the 0 to 10 Å shell and calculated the MD-minus-PDB difference at each temperature (Fig. 5C). Across all temperatures, the 7Z1X series shows consistently higher percentages of highly frustrated pairs in MD ensembles relative to PDB structures, whereas the 2P4X series exhibits no change in frustration patterns across all temperatures. Examining the 1 Å shell specifically reveals more pronounced differences in temperature sensitivity between the two series (Fig. 5D). For the 7Z1X series, the frustration difference between MD and PDB is significant across all temperatures. Near room temperatures (295 K and 300 K), this difference adopts moderate values, becoming larger at lower temperatures and smaller at higher temperatures. This temperature-dependent frustration difference closely mirrors the ΔRMSF sensitivity to temperature observed in the 7Z1X series, confirming that local frustration is the underlying determinant of interfacial dynamics. For the 2P4X series, the frustration pattern in the 1 Å shell remains identical between MD and PDB across all temperatures.

Overall, our data suggest that the local frustration pattern around the PPI interface determines interfacial dynamics. In the dynamic binding paradigm, MD sampling allows the complex to escape local minima and sample a more averaged distribution, reflected in the increased frustration observed upon thermal sampling. In contrast, the 2P4X series exhibits minimal interfacial frustration changes across all temperatures, confirming that the crystal structure serves as an appropriate proxy for the binding ensemble in this static paradigm.

### 3.5 Interface hotspot distribution in co-crystal structures confirms case-dependent role of dynamics at the residue level

In PPI engineering, such as improving binding affinity or designing interaction disruptors, identifying hotspot residues has substantial practical value. Hotspot residues are defined as those that contribute disproportionately to the binding free energy [42], and were typically assessed by alanine scanning mutagenesis. To validate the two distinct binding paradigms at the residue interaction level and relate our findings to practical nanobody engineering, we analyzed the distribution of energetically critical hotspot residues in the nanobody-antigen interfaces. Since nanobodies within each dataset bind to the same epitope, this allows us to select fixed positions of interfacial residues for alanine scanning mutagenesis (Fig. 6A) via Rosetta. We showed the ΔΔG heatmap (mutant minus wild-type) across all the datapointS within each nanobody series (Fig. 6B). Both series contained stabilizing and destabilizing mutations, but the 2P4X series exhibited more destabilizing mutations than the 7Z1X series. Using *|*ΔΔ*G| >*= 1 kcal/mol as a threshold for hotspot residues, Fig. 6C shows that in the 7Z1X series, 41.2% classified as hotspot residues. In contrast, the 2P4X series showed 75.0% as hotspot residues with only 25% classified as neutral. Furthermore, the magnitude of ΔΔG values was larger in the 2P4X series, with an average ΔΔ*G* per residue of 1.50 kcal/mol compared to 0.75 kcal/mol for the 7Z1X series. This indicates that not only does the 2P4X series have a higher percentage of hotspot residues, but the energetic penalty for disrupting these residues is also greater. Additionally, the 7Z1X dataset showed higher variability in residue contributions, with an average standard deviation of ΔΔ*G* values of 1.85 kcal/mol compared to 1.45 kcal/mol for the 2P4X series. This greater variability suggests more context-dependent roles for interfacial residues in the dynamic binding paradigm. The differential prevalence and magnitude of interface hotspots provides a residue-level rationale for our computational and dynamic findings, confirming that the two nanobody series employ fundamentally different binding strategies.

**Figure 6:**
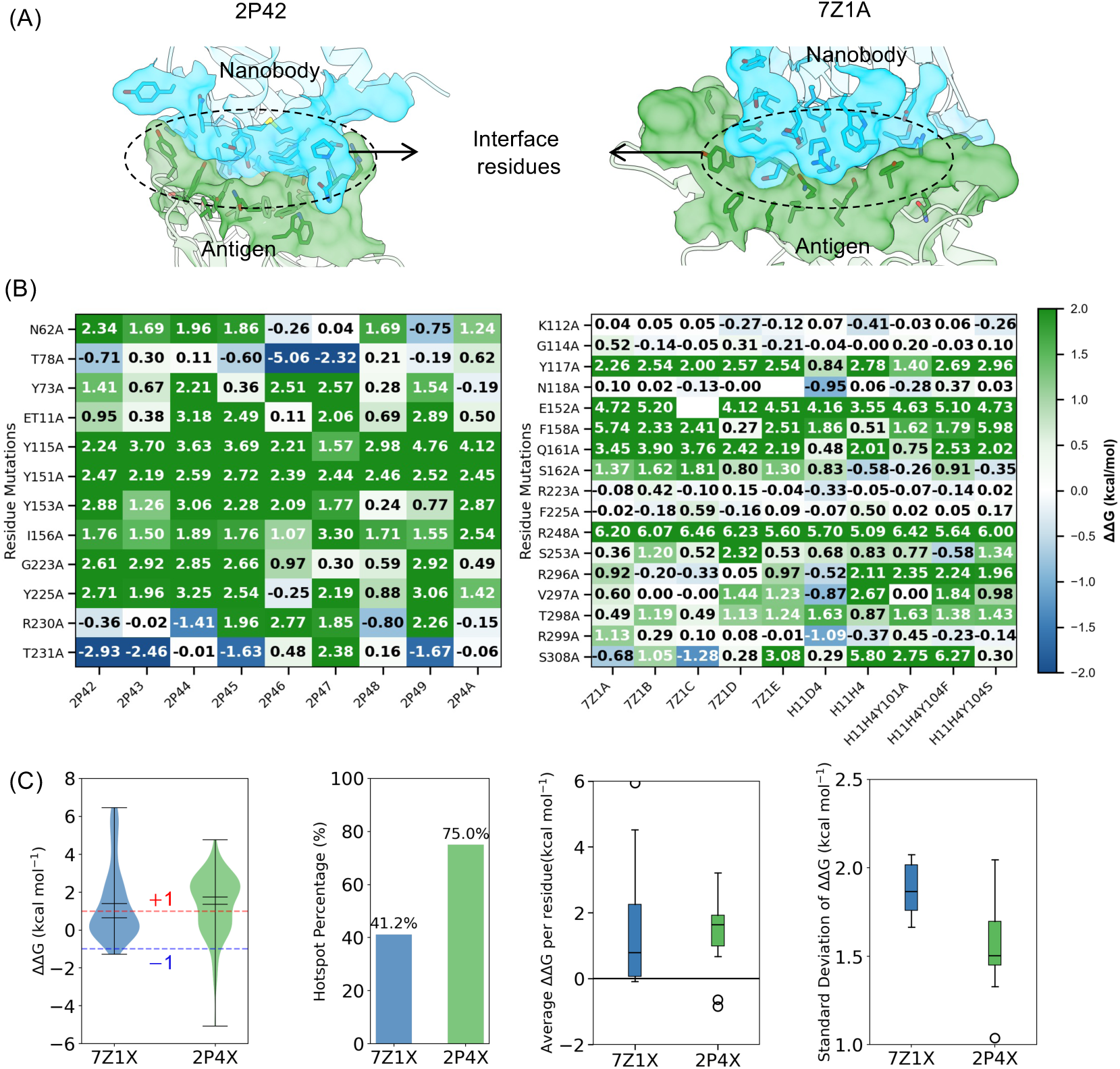
Hotspot residue distributions in the co-crystal nanobody-antigen interfaces. (A) Selection of fixed positions of interfacial residues subject to alanine scanning via Rosetta. (B) Heatmap showing the Rosetta-scored ΔΔG values (mutant minus WT) across the co-crystal structures in each dataset. (C) Statistics of ΔΔG values for the two datasets.

## 4 Discussion

While it is generally recognized that biomolecular interactions are governed by funneled energy landscapes [2], the resolution required to characterize the funnel bottom is not universal. Previous studies have noted that the critical resolution needed to reveal a binding funnel’s existence varies between protein-ligand and protein-protein systems [43]. From a structure-based perspective, we typically assume that a well-resolved co-crystal structure represents the energy minimum that determines binding affinity. However, this assumption holds only when the funnel bottom is essentially a single, deep well. If the funnel bottom is instead broad or rugged, a co-crystal static structure provides only a partial snapshot of the true thermodynamic state. Our findings demonstrate that the binding funnel bottom varies across PPI systems. In the 2P4X series, the strong performance of Rosetta scoring based on static structures indicates an optimized interface where structural complementarity drives affinity; here, the co-crystal structure resolves the funnel bottom as a smooth, deep well. In contrast, the failure of static scoring in the 7Z1X series reveals a limitation of co-crystal structures: for these complexes, only by incorporating thermal averaging from molecular dynamics simulations could we recover the affinity signal.

We further identify interfacial dynamics as the key determinant in the dynamic binding paradigm. Using MD, we show that interfacial dynamics represent a critical hidden dimension in structurally convergent binding modes. To confirm the affinity-determining role of these dynamics, we demonstrate that they are temperature-sensitive, with an optimum near physiological temperature (300 K) where they are tempered to accurately reflect experimental affinities. Both lower and higher temperatures distorted the interfacial dynamics that obscures the affinity signal. In contrast, the static paradigm exhibits negligible interfacial dynamics that are insensitive to temperature, consistent with an enthalpy-driven, lock-and-key binding mechanism. Thus, for dynamic systems, the crystal structure represents a quenched state that fails to capture the microstates required for affinity determination.

To provide a mechanistic basis for these observations, we employed local frustration theory [40]. Notably, Peter Wolynes and colleagues have pioneered energy landscapes and local frustration analysis on proteins, primarily using static PDB structures, which has fundamentally shaped our understanding of protein folding, function, and interactions [39, 40, 41]. Our use of local frustration extends the application of this theory in two distinct ways. First, we performed temperature-dependent frustration analysis using MD trajectories sampled at different temperatures to trace how interfacial frustration evolves with thermal fluctuations. Second, we focused specifically on the radial distribution function of highly frustrated pairs around the PPI interface center, which allowed us to probe the local energetic topography at the binding interface with unprecedented spatial resolution. We established a direct link between interfacial dynamics and the topology of the binding funnel. Specifically, in the static 2P4X series, the interface exhibits a lower percentage of highly frustrated pairs compared to the dynamic 7Z1X series. This minimally frustrated interface renders the complex resilient to thermal fluctuations and maintains the frustration pattern observed in the co-crystal structure across MD simulations at various temperatures. In contrast, for the 7Z1X series, MD sampling captures interfacial microstates that are suppressed in the crystal structure. These MD-revealed dynamics originate from a highly frustrated interface that creates a rugged landscape, allowing the binding partners to sample multiple microstates. This relative motion, enabled by interfacial dynamics, allows the two partners to escape kinetic traps and access the functional microstates required for accurate affinity ranking. Thus, our local frustration results unify static and dynamic paradigms within a single framework distinguished by the degree of funnel bottom ruggedness (Fig. 7).

**Figure 7:**
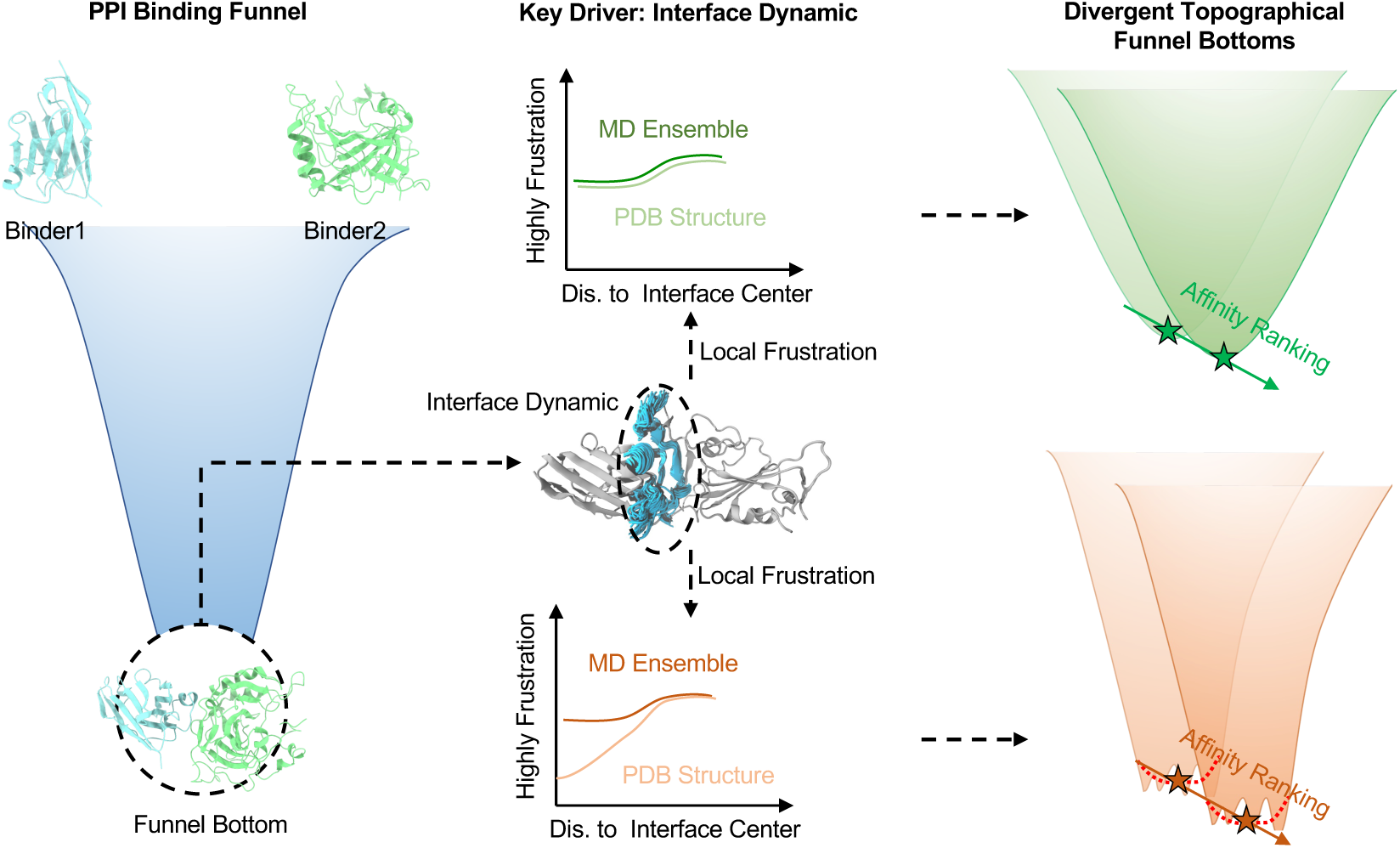
Interfacial dynamics of PPI reflects the ruggedness of the binding funnel bottom. While proteins follow a general funnel toward binding, the topography of the funnel bottom diverges based on interfacial dynamics. Similarity between static (PDB) and dynamic (MD) local frustration profiles indicates a smooth static binding paradigm (green). Conversely, divergence in these profiles reveals a highly frustrated interface, resulting in a rugged dynamic binding paradigm (orange) where affinity depends on conformational ensembles.

While we acknowledge that our major findings are based on only two datasets, these two datasets possess unique features that are particularly valuable for demonstrating that PPI binding can exist in both static and dynamic paradigms. Retrospective consultation with the original papers reporting these two nanobody series—the 2P4X series [38] and the 7Z1X series [11]—reveals that the authors had already provided experimental clues corroborating the static and dynamic binding natures, respectively. For the 2P4X series, the nanobodies are described as “well matched” to the antigen RNase A, as their binding is tolerant to dramatically different crystal packing environments, enabling the existence of six different crystal lattices [38]. This tolerance suggests a highly stable, enthalpy-driven interface with a sticky binding surface consistent with a static binding paradigm. For the dynamic 7Z1X series, Mikolajek and colleagues demonstrated using electron-density maps that global flexibility of the nanobody-antigen complex correlates with affinity [11]. This correlation directly supports the dynamic nature of this binding paradigm, where conformational sampling contributes to affinity determination. Together, these experimental observations validate that the 2P4X and 7Z1X series are representative of static and dynamic PPI binding paradigms, respectively. Furthermore, both datasets feature nanobodies that adopt nearly identical binding poses toward their cognate antigens, with affinities measured by the same research group using consistent protocols. These unique features allowed us to perform a controlled assessment of how conformational dynamics influence affinity ranking while canceling other confounding factors. The identification of such uniquely well-controlled nanobody series was therefore instrumental in enabling us to demonstrate that PPI binding can exist in both dynamic and static paradigms. Although the two sets are relatively small, their unique features allowed us to perform a controlled assessment of how conformational dynamics influence affinity ranking while canceling other confounding factors, such as epitope differences.

From a practical perspective, a major challenge in computational biophysics is determining a priori whether a system requires ensemble-based sampling. Our findings indicate that local frustration patterns provide a principled metric for this determination. To demonstrate the generalizability of our findings, we identified two additional nanobodies, Nb1 and Nb6, that bind to the same EGF protein via entropy– and enthalpy-driven binding mechanisms, respectively [9]. Applying the conceptual framework (summarized in Figure 7) to Nb1 and Nb6 successfully identified their dynamic and static binding natures (see Fig. S11 for details). Specifically, the entropy-driven Nb1 exhibited a significant difference between single static structure based and MD-based local frustration patterns at the interface, while the enthalpy-driven Nb6 showed nearly identical patterns—consistent with our classification framework. These encouraging results demonstrate that our findings, based on the 2P4X and 7Z1X series, are applicable to additional nanobody PPIs. More importantly, these new results suggest that our proposed method for distinguishing dynamic and static PPIs based on interfacial frustration analysis holds practical application potential. By classifying a complex as static or dynamic through frustration analysis, one can select the appropriate modeling strategy—static structure-based scoring or ensemble-based sampling—thereby balancing predictive accuracy with computational cost. Our proposed framework advances beyond uniform prediction methods toward a landscape-informed approach for PPI analysis and design.

## 5 Conclusions

In this study, we combined MD simulations, ensemble-based scoring, and local frustration analysis to investigate how conformational dynamics contribute to PPI binding affinity. Using nanobody-antigen systems as models, we demonstrated that PPI binding paradigms can have fundamentally different energy landscape topographies. In the static paradigm (2P4X series), the binding funnel bottom is smooth and minimally frustrated, and co-crystal structures alone can correctly rank experimental affinities within this series. Here, structural complementarity encodes the thermodynamic information, and conformational sampling introduces only noise. In the dynamic paradigm (7Z1X series), the funnel bottom is rugged and highly frustrated, requiring ensemble-based sampling at the appropriate temperature to recover the affinity signal. Interfacial dynamics in this series are temperature-sensitive, with an optimum near 298 K where the sampled microstates correctly reflect experimental affinity rankings. Local frustration analysis revealed that these paradigms are distinguished by the degree of interfacial ruggedness. Dynamic interfaces exhibit a higher density of frustrated interactions that drive conformational sampling, whereas static interfaces rely on a smaller set of optimized hotspot residues. This frustration-based framework explains how structurally convergent binding modes can encode diverse affinities through interfacial plasticity, suggesting that structural identity does not imply dynamic identity. Moreover, we propose the local frustration topography of the interface as a practical method for quickly assessing whether a PPI belongs to the static or dynamic paradigm, and we validated its applicability on additional nanobodies beyond the two primary series. Our work provides a principled metric for diagnosing binding paradigms and advances toward landscape-informed affinity prediction in protein engineering and therapeutic design.

## 6 Acknowledgments

Research reported in this work was supported by the Natural Science Foundation of China (32501103), Natural Science Foundation of Heilongjiang Province (PL2024B022), China Postdoctoral Science Foundation (2023M730827, 2025MD784111), Heilongjiang Provincial Postdoctoral Science Foundation (LBH-Z24211).

